# Low frequency stimulation for seizure suppression: identification of optimal targets in the entorhinal-hippocampal circuit

**DOI:** 10.1101/2024.02.09.579614

**Authors:** Piret Kleis, Enya Paschen, Ute Häussler, Carola A. Haas

## Abstract

**Background:** Mesial temporal lobe epilepsy (MTLE) with hippocampal sclerosis (HS) is a common form of drug-resistant focal epilepsy in adults. Treatment for pharmacoresistant patients remains a challenge, with deep brain stimulation (DBS) showing promise for alleviating intractable seizures. This study explores the efficacy of low-frequency stimulation (LFS) on specific neuronal targets within the entorhinal-hippocampal circuit in a mouse model of MTLE.

**Objective/Hypothesis:** Our previous research demonstrated that LFS of the medial perforant path (MPP) fibers in the sclerotic hippocampus reduced seizures in epileptic mice. Here, we aimed to identify the critical neuronal population responsible for this antiepileptic effect by optogenetically stimulating presynaptic and postsynaptic compartments of the MPP-dentate granule cell (DGC) synapse at 1 Hz. We hypothesize that specific targets for LFS can differentially influence seizure activity depending on the cellular identity and location within or outside the seizure focus.

**Methods:** We utilized the intrahippocampal kainate (ihKA) mouse model of MTLE and targeted specific neural populations using Channelrhodopsin2 (ChR2) and stereotactic optic fiber implantation. We recorded intracranial neuronal activity from freely moving chronically epileptic mice with and without optogenetic LFS up to three hours.

**Results:** We found that LFS of MPP fibers in the sclerotic hippocampus effectively suppressed epileptiform activity while stimulating principal cells in the MEC had no impact. Targeting DGCs in the sclerotic septal or non-sclerotic temporal hippocampus with LFS did not reduce seizure numbers but shortened the epileptiform bursts.

**Conclusion:** Presynaptic stimulation of the MPP-DGC synapse within the sclerotic hippocampus is critical for seizure suppression via LFS.

## Introduction

MTLE is the most common form of drug-resistant focal epilepsy in adults, with recurrent seizures originating from hippocampal or surrounding mesiotemporal structures [1,2]. The main histological hallmark associated with refractory MTLE is HS, characterized by neuronal loss and reactive gliosis [3]. Pharmacoresistant patients rely on invasive treatments such as resective surgery or DBS for seizure control. DBS offers advantages like non-lesional nature, reversibility, and adjustability [4]. However, if the seizure focus is unilateral, clearly identifiable, and not in a region of predictable cognitive risk, then resection is still preferred over DBS due to better seizure freedom outcomes [4–6]. Thus, neurostimulation approaches could be improved by optimizing target areas and stimulation parameters for specific epilepsy types.

Epilepsy patients eligible for DBS typically undergo high-frequency stimulation (HFS, >100Hz) in the seizure focus or a thalamic nucleus (anterior or centromedial) to impede seizure propagation to other brain regions [7,8]. The outcome of DBS in MTLE patients with HS is controversially reported: some studies show a worse response to hippocampal DBS compared to non-lesional MTLE [9,10], while others find the opposite [5]. Alternatively, LFS (<10 Hz) targeting the hippocampus directly [11] or via the fornix [12,13] exerted antiepileptic effects also in the presence of HS. In addition, a recent study showed that closed-loop responsive neurostimulation, which conventionally involves applying HFS upon seizure detection, can be switched from HFS to LFS in non-responsive cases, yielding significant seizure reduction [14]. Given the heterogeneity of the clinical cohorts and the dependence on DBS parameters and epilepsy type, studies in animal models are needed to investigate stimulation strategies and to understand the contribution of different cell types to the antiepileptic effect of LFS.

Cell-specific optogenetic methods have been pivotal in epilepsy research to investigate the mechanisms of seizure initiation, propagation, and termination and to identify cell populations as therapeutic targets [15–17]. Focal seizures are believed to result from excitation-inhibition imbalances [18–20]. Thus, most optogenetic interventions in models of focal epilepsy target this imbalance by activating interneurons using cation-permeable channelrhodopsin 2 (ChR2) [21–23] or silencing principal cells with anion-, proton-, or potassium-conducting optogenetic tools [21,24–26]. The optimal stimulation target, in our perspective, is a cell population that facilitates seizure control through activation rather than inhibition for clinical translatability via electric stimulation.

In the present study, we modeled MTLE with HS using ihKA mice. After kainate (KA)-induced *status epilepticus* (SE), mice develop chronic epilepsy within 2-3 weeks with recurrent focal seizures arising from the KA-injected sclerotic hippocampus, considered as seizure focus [27–32]. Similarly to MTLE patients with HS type I, ihKA mice show severe loss of pyramidal neurons in CA1 and CA3 regions and extensive loss of interneurons across the sclerotic hippocampus [3,33–35]. The cell populations preserved in the sclerotic hippocampus, presenting potential stimulation targets, are CA2 pyramidal cells, DGCs, and incoming projections such as MPP fibers from the medial entorhinal cortex (MEC) [27,35–37]. The neuronal loss decreases from septal towards temporal hippocampus, thus cell populations in the temporal hippocampus are also potential targets for LFS [32].

The dentate gate hypothesis posits that the MPP-DGC connection is the gateway to the hippocampus, and the breakdown of the gate leads to over-excitation and seizures [38–40]. DGCs in the sclerotic hippocampus have severely impaired intrinsic properties [41,42] that contribute to the dysfunction of the dentate gate function [41–43]. Furthermore, the MPP-DGC synapse shows signs of long-term potentiation-associated ultrastructural changes such as enlarged pre- and postsynaptic compartments and increased synapse number [35]. Therefore, we targeted the MPP-DGC synapse with LFS and showed that 1 Hz optogenetic LFS of the MPP fibers in the sclerotic hippocampus suppresses spontaneous focal seizures and prevents induced behavioral seizures [44].

In this study, we selectively modulate the presynaptic and postsynaptic sites of the dentate gate to further investigate the seizure-suppressive effects of hippocampal LFS. Furthermore, we aim to elucidate whether the optimal target for LFS is in the sclerotic hippocampus or in its vicinity. Thus, we probe the antiepileptic effect of LFS in four cellular targets: (1) MPP fibers in the sclerotic hippocampus, (2) principal cells in the MEC, (3) DGCs in the sclerotic hippocampus, and (4) DGCs in the non-sclerotic temporal hippocampus. We present evidence that LFS of DGCs shortens epileptiform events, while reliable seizure suppression is attained only with MPP stimulation within the seizure focus.

## Material and methods

### Animals

Experiments were performed in adult transgenic male mice. For targeting entorhinal principal cells, we used C57BL/6 Tg(Thy1-eGFP)-M-Line [45]. For specific targeting of DGCs, we crossed C57BL/6-Tg(Rbp4-cre)KL100Gsat mice with Ai32(RCL-ChR2(H134R)/EYFP mice resulting in transgenic mice expressing ChR2 in retinol binding protein 4(Rbp4)-positive cells (Rbp4-Cre-Ai32 mice). A total of 40 mice bred at CEMT Freiburg were used specifically for this study. We reduced the number of animals by reevaluating data from a previous study, and all procedures carried out with these mice are described in detail in [44]. Mice were kept in a 12 h light/dark cycle at room temperature with food and water *ad libitum*. All animal procedures were carried out following the guidelines of the European Community’s Council Directive of 22 September 2010 (2010/63/EU) and were approved by the regional council (Regierungspräsidium Freiburg).

### KA and virus injections

Mice were injected with 50 nl KA (10 or 15 mM, Tocris) into the right dorsal hippocampus, as described previously [35,44]. To reduce mortality rates, the KA concentration was reduced from 15 to 10 mM during the study. The stereotaxic injection was performed under deep anesthesia (ketamine hydrochloride 100 mg/kg, xylazine 5 mg/kg, atropine 0.1 mg/kg body weight, i.p.) using a Nanoject III (Drummond Scientific Company). Stereotaxic coordinates (in mm) for ihKA were anterioposterior (AP) −2.0, mediolateral (ML) −1.5 relative to bregma, and dorsoventral (DV) −1.5 relative to the cortical surface. Mice in which Ca^2+^/calmodulin-dependent protein kinase II α (CaMKIIα) expressing entorhinal principal cells were optogenetically targeted, received an additional recombinant adeno-associated virus (AAV1.CaMKIIα.hChR2(H134R)-mCherry.WPRE.hGH, 450 nl, Viral Vector Facility University of Zurich) injection into the MEC (AP −5.0, ML −2.9, DV −1.8) during the same surgery. Following the ihKA injection, behavioral SE was verified by observation of mild convulsions, immobility, or rotations, as described before [28,46]. Table A1 summarizes sample sizes and reasons for exclusion.

### Electrode and optic fiber implantations

Teflon-coated platinum-iridium wires (125 µm diameter; World Precision Instruments) were implanted 14-17 days after SE into ipsilateral and contralateral hippocampus at the level of KA injections (iHC and cHC, respectively) for local field potential (LFP) recordings. Stereotaxic coordinates of the recording electrodes were (in mm) AP −2.0, ML +1.4 (cHC) or −1.4 (iHC) relative to bregma, and DV −1.6 relative to the cortical surface. An optic fiber was implanted into one of the target regions in the ipsilateral hemisphere, either: (1) MEC together with an additional recording electrode at an 8° angle (from transverse sinus: AP +0.2, from bregma: ML −2.9, from cortical surface: −2.3, in mm), (2) septal hippocampus next to the iHC electrode at a 30° angle (AP −2.0, ML −2.4, DV −1.0, in mm), or (3) temporal hippocampus (tHC) together with an additional recording electrode (AP −3.4, ML −2.8, DV −2.1, in mm).

Two stainless steel screws (DIN 84) were implanted above the frontal cortex to provide a reference and ground. Electrodes and screws were soldered to a micro-connector (BLR1-type) and fixed with dental cement (Paladur). The electrode and optic fiber positions were verified by post hoc histology as described previously [44].

### Electrophysiological recordings and optogenetic stimulation

Local field potentials (LFPs) were recorded from freely moving mice in the chronic phase of epilepsy between 30-40 days after ihKA injections. Mice were connected to a miniature preamplifier (MPA8i, Smart Ephys/Multi Channel Systems) and signals were amplified 1000-fold, bandpass-filtered from 1 Hz to 5 kHz and digitized with a sampling rate of 10 kHz (Power1401 analog-to-digital converter, Spike2 software, Cambridge Electronic Design).

First, we acquired a three-hour-long baseline recording without stimulation. Subsequently, the mice were optogenetically stimulated with pulsed blue light (460 nm; 50 ms pulses; 150 mW/mm^2,^ blue LED from Prizmatix) at 1 Hz and simultaneously LFP recorded for three hours on two separate days. The MPP stimulation protocol has been described previously [44].

### Tissue preparation and immunohistochemistry

At the end of each experiment, mice were deeply anesthetized and transcardially perfused with 0.9% saline followed by 4% paraformaldehyde in 0.1 M phosphate buffer (PB, pH 7.4). Dissected brains were post-fixed overnight, transferred to PB, and sectioned (coronal plane, 50 μm) with a vibratome (VT100S, Leica Biosystems). Slices were collected and stored in PB for immunohistochemistry.

To determine optic fiber and electrode positions and to validate hippocampal sclerosis, we performed immunohistochemistry with markers for neurons (NeuN) and astrocytes (glial fibrillary acidic protein, GFAP). Free-floating sections were pre-treated with 0.25% TritonX-100 and 1% bovine serum albumin (Sigma-Aldrich) diluted in PB for 30 minutes. Subsequently, slices were incubated with guinea-pig anti-NeuN (1:1000; Synaptic Systems) and rabbit anti-GFAP (1:500, Dako) overnight at 4°C. Sections were rinsed and then incubated for 2.5 h in donkey anti-guinea pig Cy5- and goat anti-rabbit Cy3-conjugated secondary antibody (1:200, Jackson ImmunoResearch Laboratories Inc.) followed by extensive rinsing in PB. The sections were mounted on glass slides and coverslipped with Immu-Mount^TM^ medium (Thermo Shandon Ltd).

### Image acquisition and histology

Tiled fluorescent images of the brain sections were taken with an *AxioImager 2* microscope (Carl Zeiss Microscopy GmbH) using a Plan-Apochromat 10x objective with a numerical aperture of 0.45 (Zeiss). The presence of unilateral HS in the epileptic mice was confirmed in NeuN-labeled sections showing DGC dispersion, cell loss in CA1 and CA3, and in GFAP-labeled sections showing astrogliosis (Fig. A1).

### Analysis of epileptiform activity

Downsampled (to 500 Hz) hippocampal LFPs recorded from epileptic animals were analyzed using a custom-made semi-automated algorithm that detects and classifies epileptiform activity [47]. In the ihKA mouse model, epileptiform activity occurs as single sharp wave epileptiform spikes and as bursts, which are clusters of many spikes [28]. The algorithm classifies the epileptiform bursts according to their spike-load into low-load (LL), medium-load (ML), and high-load bursts (HL) with the latter corresponding to electrographic focal seizures [47]. To assess the effect of LFS at different targets on seizure activity, we calculated the ‘HL burst ratio’, which is the fraction of time spent in HL bursts from the total recording time. Furthermore, we compared the HL burst number and the cumulative distribution of epileptiform burst durations, including all the burst classes, in baseline and LFS recordings. Generalized seizures and the following 30 min of the LFP recording were excluded from the analysis due to long-lasting suppression of neuronal activity after such a seizure.

### Statistical analysis

Data were tested for significant differences with Prism 9 software (GraphPad Software Inc.). A paired t-test was used to compare two matched groups of parametric data. Comparisons of more than two parametric data sets were performed either with a one-way ANOVA or with repeated measures (RM) ANOVA, in the case of matched groups. If an ANOVA indicated that not all group means were equal, Tukey’s multiple comparisons test was performed additionally. Cumulative distribution plots were compared using the Kruskal-Wallis test. Significance thresholds were set to: *p<0.05, **p<0.01. For parametric data, mean and standard error of the mean (SEM) are reported.

## Results

### LFS of MPP fibers in the sclerotic hippocampus suppresses spontaneous seizures

In this study, we first reevaluated a dataset of optogenetic MPP stimulation in the sclerotic hippocampus from Paschen et al., (2020) [44] in order to compare it to targets inside and outside the sclerotic seizure focus. This includes data recorded with an electrode implanted side-by-side with an optic fiber into the molecular layer of the dentate gyrus at the level of KA injection (Fig. 1A, B). During the chronic phase of epilepsy (>21 days after KA), four-hour LFP recordings were acquired in freely moving mice including one-hour pre-recording, one-hour optogenetic LFS, and two hour-long post recordings (Fig. 1C). Using a semi-automated algorithm to detect and classify epileptiform activity [48] (Fig. 1D), we observed spontaneous epileptiform bursts of all classes (HL, ML, and LL) arising frequently during pre-recordings (Fig. 1D). While the MPP fibers were optogenetically stimulated at 1 Hz, the HL bursts, corresponding to focal seizures, were rare (Fig. 1E). We quantified the HL burst ratio and burst number across animals and averaged the results from two sessions. We found that the burst ratio was significantly lower during LFS (0.05 ± 0.03) compared to pre-recording (0.18 ± 0.03, RM ANOVA: F=9.64, n=6, p=0.0065, Tukey’s multiple comparison test: p=0.005; Fig. 1F). In the first hour after stimulation, the burst ratio (0.16 ± 0.05) was restored to baseline level (Tukey’s multiple comparison test: p=0.85), suggesting that there is no long-lasting effect of LFS. The average HL burst number per hour was 14.42 ± 1.90 during pre-recording and significantly decreased during LFS to 3.25 ± 1.7 (paired t-test: t =5.14, n=6, p=0.004; Fig. 1G). Furthermore, we looked into the effect of LFS on the duration of LL, ML, and HL bursts, and found a clear shift of the cumulative distribution towards shorter bursts during LFS (Kolmogorov–Smirnov test, n=6, p<0.001; Fig. 1H). All in all, optogenetic LFS of MPP fibers in the sclerotic hippocampus shortened the epileptiform bursts, strongly reduced the seizure count and the time spent in focal seizures.

**Figure 1.**
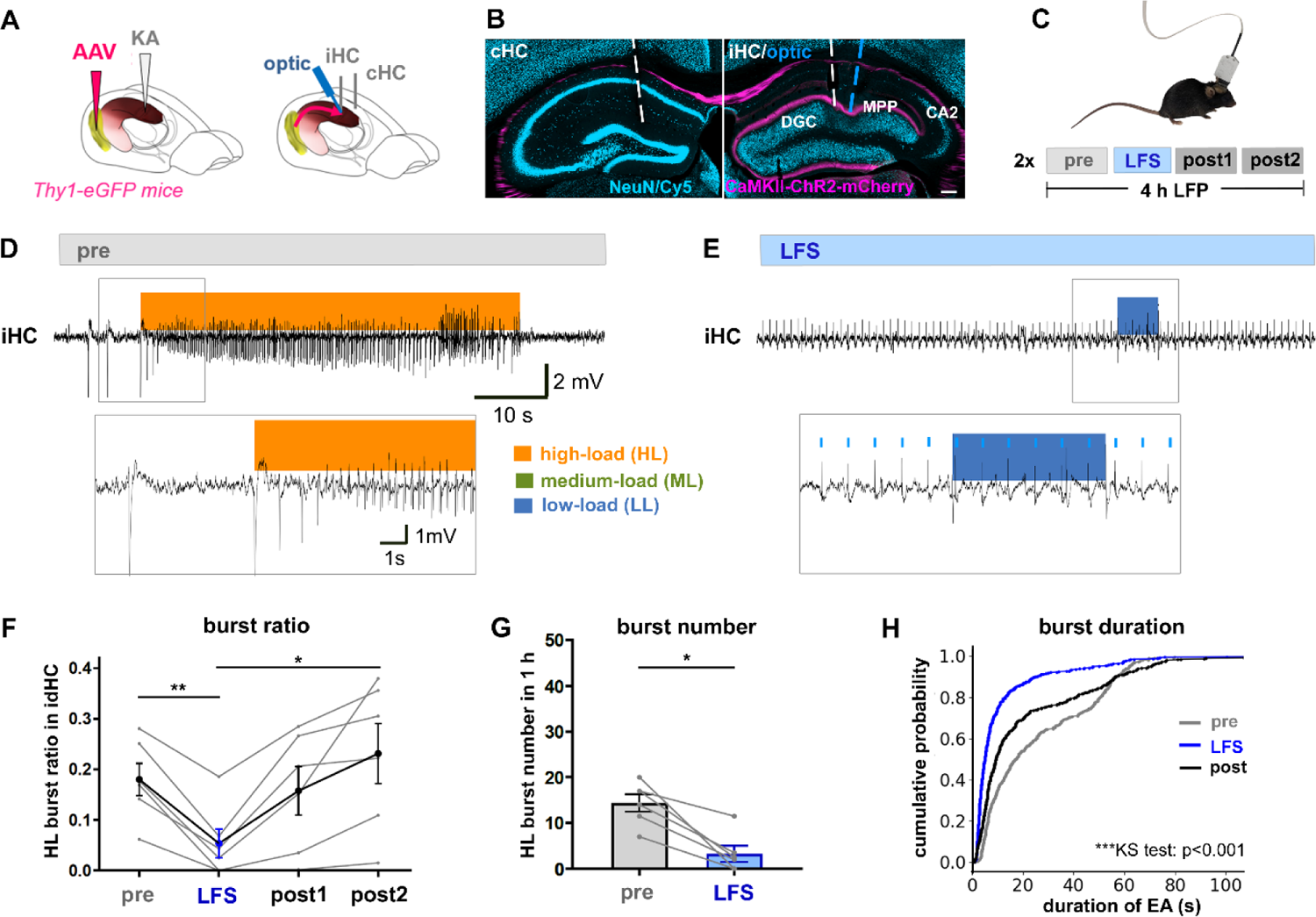
Targeting MPP fibers in the sclerotic hippocampus with LFS. (A) Injection and implantation scheme. We injected KA into the right dorsal hippocampus and the AAV carrying ChR2 to CaMKIIα-positive neurons into the right MEC. The electrodes were implanted into each hippocampus, the optic fiber tip targeted the MPP fibers in the right dentate gyrus. (B) Histological validation of ChR2-mCherry expression and electrode/optic fiber positions in the dentate gyrus. Scale bar 200 µm. (C) A session consisted of one-hour-long baseline (pre) recording, followed by one-hour LFS phase with 1 Hz light pulses and two hour-long post-recordings (post1, post2). Each mouse had two sessions on separate days. (D) A representative LFP trace from a pre-recording showing a typical seizure-like event, i.e. HL burst detected in the iHC. (E) A representative LFP during optogenetic LFS of MPP fibers in the sclerotic hippocampus demonstrates robust evoked responses to light pulses and no HL burst activity. (F) HL burst ratio was significantly reduced during the LFS but restored to the pre-stimulation level after the stimulation was turned off, RM ANOVA with post-hoc test **p<0.01, *p<0.05, n=6 (the average of two sessions per mouse as grey lines, the overall mean as black line). (G) The number of HL bursts was significantly lower during the LFS compared to pre-recording, paired t-test *p<0.05, n=6 (average of two sessions per mouse as grey lines). Mean with SEM presented as error bars. (H) The cumulative distribution function of epileptiform activity (EA) durations during pre-, LFS- and post1-recordings. Duration of epileptiform bursts was significantly reduced during LFS compared to pre-recordings Kolmogorov–Smirnov test, ***<0.001.

### LFS of principal cells in the MEC has no effect on epileptiform activity

Similar to MPP fiber stimulation in the sclerotic hippocampus, the presynapse of the dentate gate can also be stimulated outside the seizure focus by direct stimulation of principal cells in the MEC. Hence, we used a similar experimental scheme as before but inserted the optic fiber with a recording electrode into the MEC (Fig. 2A, B). We first acquired three-hour LFP recordings to assess the baseline epileptiform activity, then applied optogenetic LFS of CaMKIIα-positive cells in the MEC for three hours on two following days. HL bursts were present in both iHC and MEC (Fig. 2D), however in the MEC, the amplitude of epileptiform activity was low, and sparse interictal spikes were observed more often than epileptiform bursts. During LFS, responses to light pulses were evident in MEC but the propagation to the ipsilateral hippocampus was unreliable (Fig. 2E). We quantified the epileptiform bursts in the sclerotic hippocampus and found that optogenetic LFS in MEC had no effect on HL burst ratio (baseline: 0.20 ± 0.04, LFS: 0.21 ± 0.05, paired t-test: t=0.67, n=5, p=0.54, Fig 2F). Also, the average HL burst number (baseline: 15.47 ± 3.90, LFS: 17.55 ± 3.51, paired t-test: t=0.29, n=5, p=0.78, Fig. 2G) and burst duration (Kolmogorov–Smirnov test, n=5, p=0.07; Fig. 2H) were similar in baseline and LFS sessions. Therefore, targeting principal cells in the MEC with LFS did not have the same seizure-suppressive effect as the downstream MPP stimulation.

**Figure 2.**
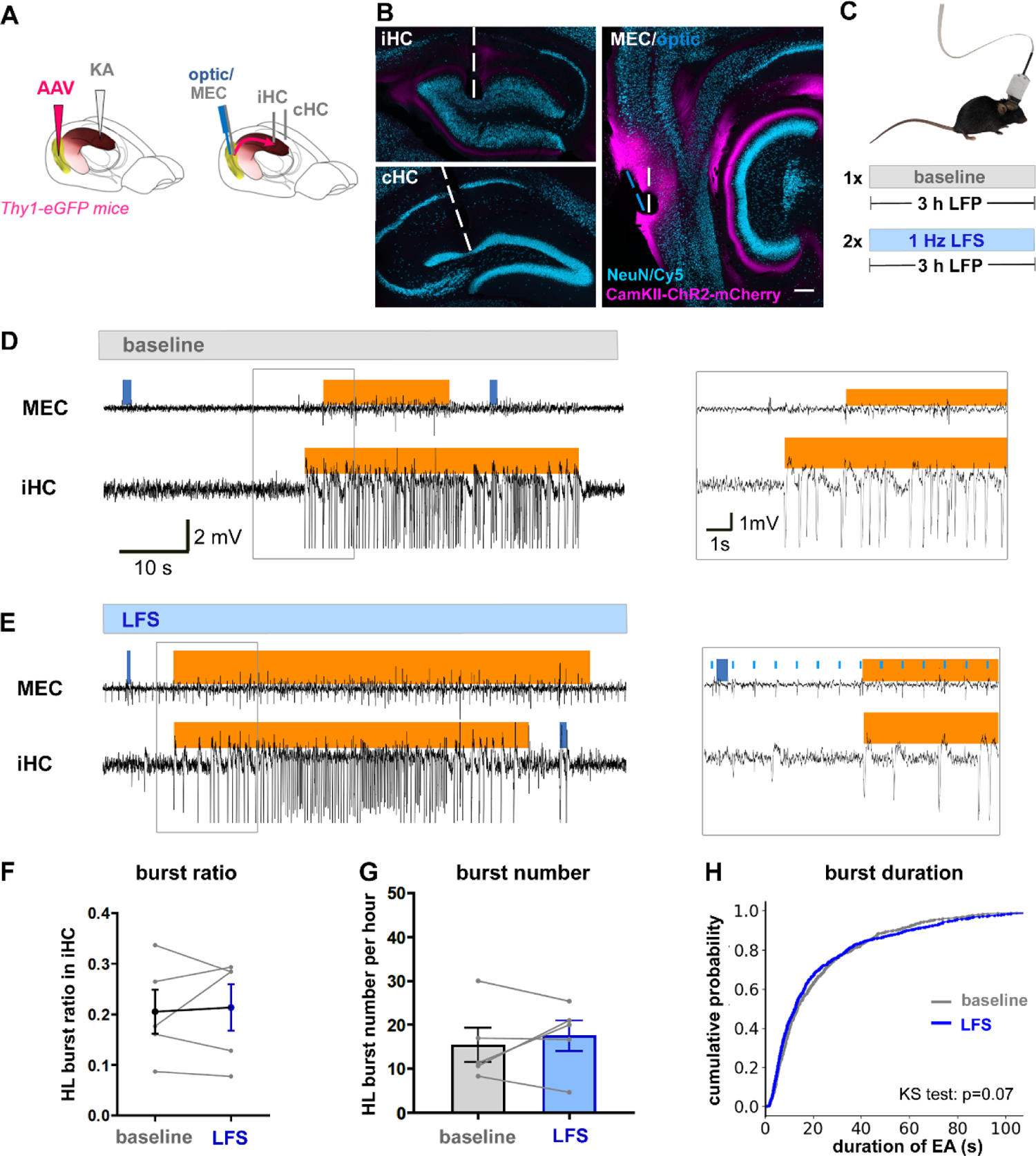
Targeting excitatory neurons in the MEC with LFS. (A) Injection and implantation scheme. IhKA-injected mice received a viral vector into the right MEC for ChR2 expression in CaMKIIα-positive neurons. Recording electrodes were implanted into each dentate gyrus (iHC, cHC) and the right MEC, the MEC electrode having the optic fiber attached in parallel. (B) Histological validation of ChR2-mCherry expression and electrode/optic fiber positions in iHC, cHC and MEC. Scale bar 200 µm. (C) A three-hour-long LFP recording without (baseline) and two with optogenetic stimulation (LFS) were performed on separate days. (D) A representative trace from a baseline LFP with a HL burst in MEC and iHC. (E) A representative LFP during optogenetic LFS of excitatory MEC neurons showing clear evoked responses to light pulses in MEC but not in iHC. (F) HL burst ratio was unaltered by the LFS, paired t-test p>0.05, n=5 (the average of two sessions per mouse as grey lines, the overall mean as black line). (G) The number of HL bursts was not significantly different during the LFS compared to baseline, paired t-test p>0.05, n=5 (grey lines). Mean with SEM presented as error bars. (H) There was no change of the cumulative distribution function comparing the burst durations during baseline and LFS (Kolmogorov–Smirnov test, p=0.07, n=5 mice).

### LFS of DGCs in the sclerotic hippocampus has a pro-epileptic effect

In contrast to the presynaptic MPP or MEC stimulation, modulation of the postsynaptic counterpart of the dentate gate can be achieved by applying LFS directly to DGCs in the sclerotic hippocampus. To this end, we used the transgenic Rbp4-cre-Ai32 mouse line that expresses ChR2-YFP exclusively in Rbp4-positive DGCs. These mice were injected with KA and implanted in the same way as the MPP stimulation group (Fig. 3A). The implantation positions and the expression of ChR2-YFP in DGC dendrites and axons were confirmed by histology (Fig. 3B). Again, we acquired three-hour baseline recordings and two recordings with optogenetic LFS (Fig. 3C-E). We found that LFS of DGCs in the sclerotic hippocampus did not alter the HL burst ratio (baseline: 0.15 ± 0.05, LFS: 0.15 ± 0.04, paired t-test: t=0.04, n=7, p=0.96; Fig. 3F) but increased the HL burst number (baseline: 13.71 ± 3.08, LFS: 25.00 ± 4.66, paired t-test: t=2.84, n=7, p=0.029; Fig. 3G). However, the bursts that occurred were shorter during LFS compared to baseline (Kolmogorov–Smirnov test: n=7, p<0.001; Fig. 3H).

**Figure 3.**
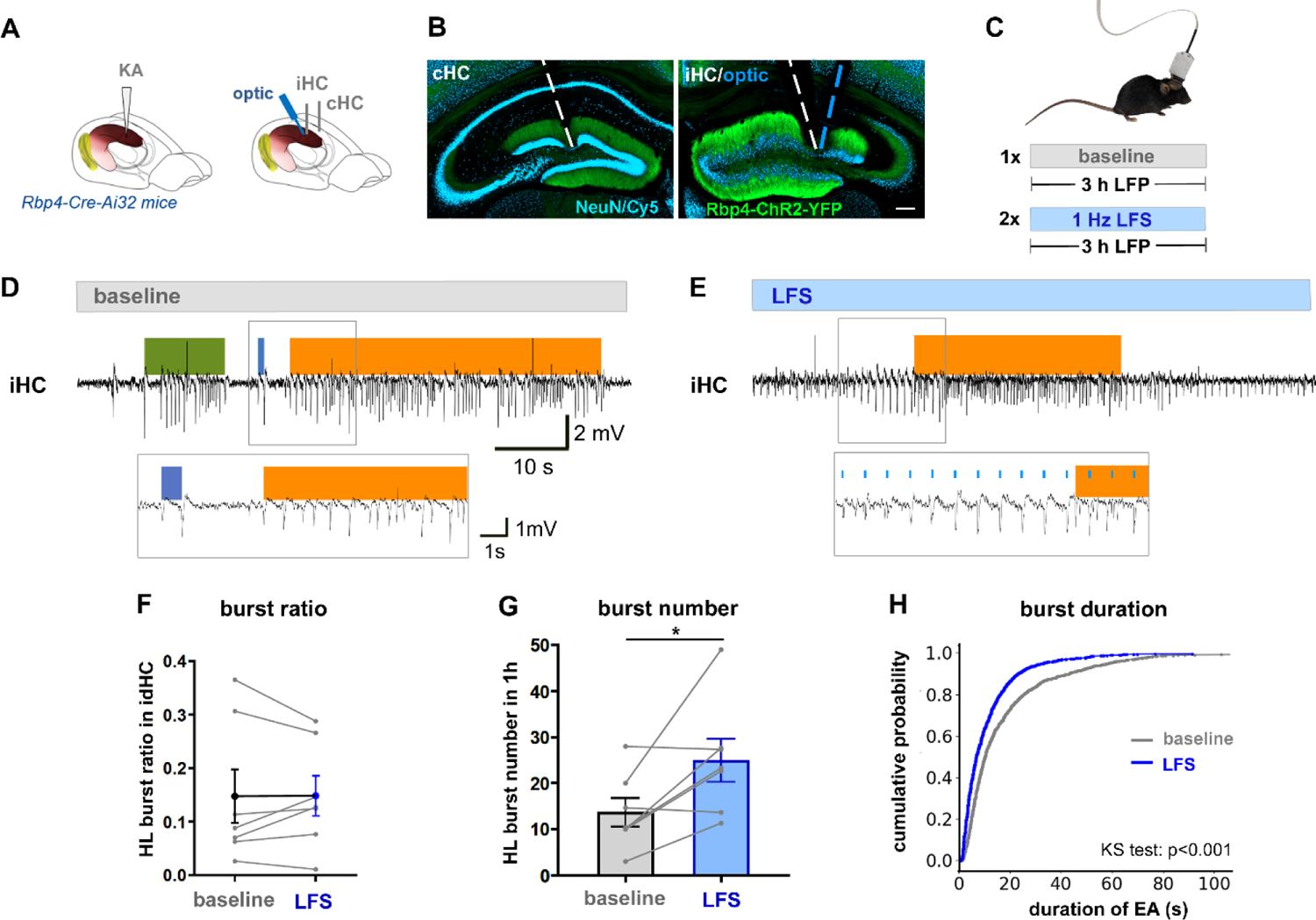
Targeting DGCs in the sclerotic hippocampus with LFS. (A) Injection and implantation scheme for Rbp4-Cre-Ai32 mice. (B) Histological validation of ChR2-YFP expression in DGC dendrites and axons, electrode/optic fiber positions in iHC and cHC. Scale bar 200 µm. (C) Three-hour-long LFP recordings without (baseline) and with LFS optogenetic stimulation were performed on separate days. (D) A representative trace from a baseline LFP with a LL, ML, and HL burst. (E) A representative LFP during LFS of DGCs showing clear evoked responses to light pulses in iHC. (F) HL burst ratio was unaltered by the LFS, paired t-test p>0.05, n=6 (the average of two sessions per mouse as grey lines, the overall mean as black line). (G) The number of HL bursts was significantly increased during the LFS compared to baseline, paired t-test p=0.03, n=6 (grey lines). Mean with SEM presented as error bars. (H) Duration of epileptiform bursts, including LL, ML, and HL, was significantly reduced during LFS compared to baseline (Kolmogorov–Smirnov test, p<0.001).

### LFS of DGCs in the temporal hippocampus shortens epileptiform bursts

DGCs in the septal sclerotic hippocampus are dysfunctional [49,50] and surrounded by a reorganized hippocampal network, including recurrent connections via DGC autapses [27], reduced inhibitory drive [33,51], and lost postsynaptic partners such as CA3 pyramidal neurons and mossy cells [27,52]. However, in the temporal non-sclerotic hippocampus of ihKA mice, the hippocampal circuitry is preserved. Thus, we hypothesized that LFS of DGCs in the non-sclerotic temporal hippocampus could have a different effect than in the sclerotic hippocampus. We again used the Rbp4-Cre-Ai32 mice for ihKA injection and implanted recording electrodes into both septal hippocampi and an optic fiber parallel to the recording electrode into the temporal hippocampus (Fig. 4A-B). During three-hour baseline recordings, epileptiform activity was detected in both the septal and temporal hippocampus (Fig. 4D) as described previously [32]. Activation of DGCs with optogenetic LFS in the temporal hippocampus evoked stable responses at the stimulation site (Fig. 4E), but the propagation to the septal hippocampus was unstable and seemed to occur more reliably around epileptiform events. LFS of DGC in the non-sclerotic temporal hippocampus neither altered the HL burst ratio (baseline: 0.16± 0.05, LFS: 0.13± 0.03, paired t-test: 0.96, n=6, p=0.38; Fig 4F), nor HL burst number (baseline: 13.90 ± 3.35, LFS: 16.29 ± 3.52, paired t-test: t=0.56, n=7, p=0.60; Fig. 4G). However, similarly to LFS of septal DGCs, LFS of DGCs in the temporal hippocampus significantly shortened the epileptiform bursts (Kolmogorov–Smirnov test: n=6, p<0.001; Fig. 4H).

**Figure 4.**
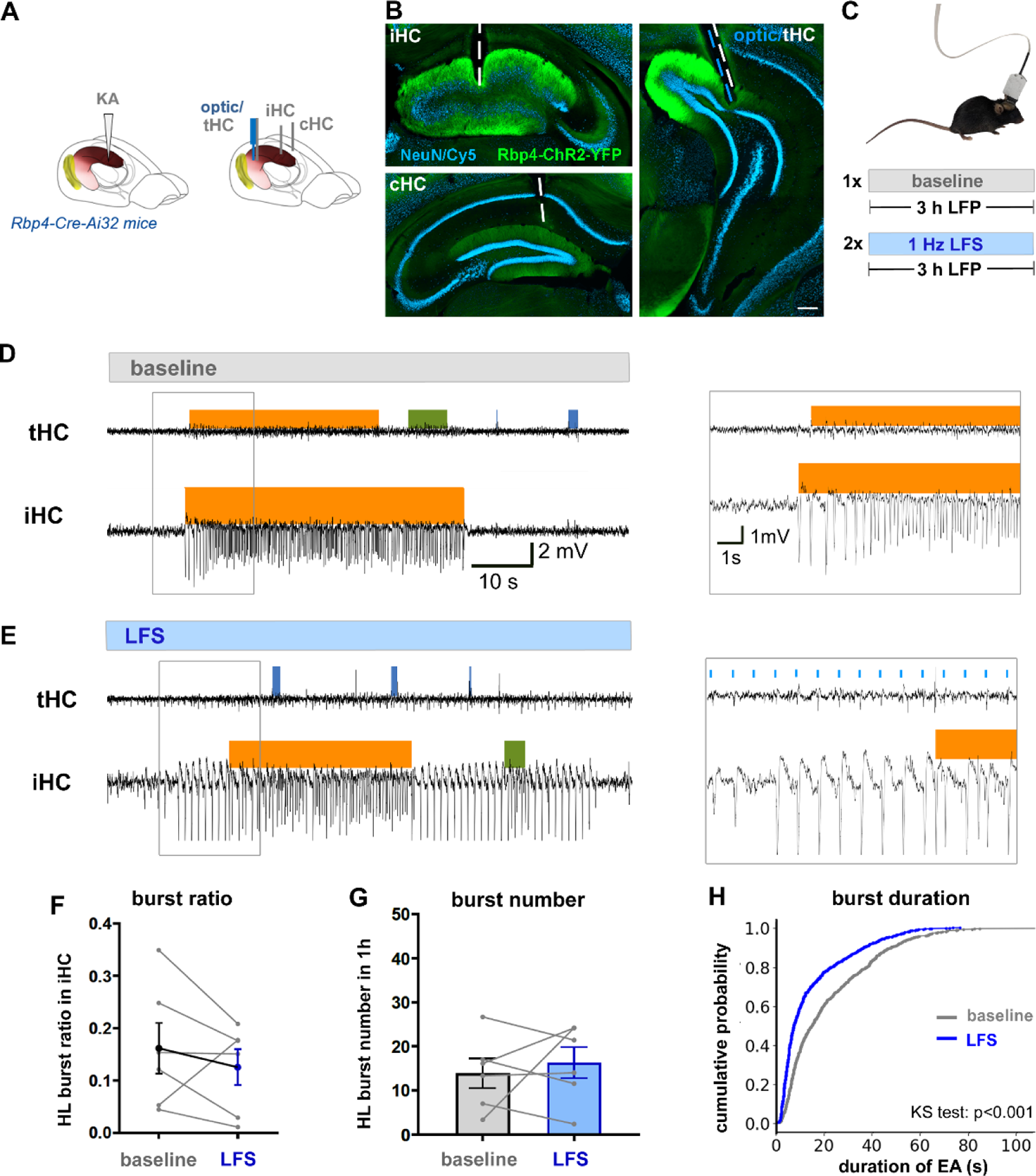
Targeting DGCs in the non-sclerotic temporal hippocampus with LFS. (A) KA injection and implantation scheme in Rbp4-Cre-Ai32 mice. Here, the optic fiber was placed together with a recording electrode into the dentate gyrus of the temporal hippocampus. (B) Histological validation of ChR2-YFP expression in DGC and electrode/optic fiber positions in iHC, cHC, and tHC. Scale bar 200 µm. (C) Three-hour-long LFP recordings without (baseline) and with (LFS) optogenetic stimulation were performed on separate days. (D) A representative trace from a baseline LFP with a LL, ML, and HL burst at both ipsilateral recording sites. (E) A representative LFP during LFS of DGCs in the temporal hippocampus showing evoked responses in tHC that sometimes propagate to iHC with large amplitudes. (F) HL burst ratio and (G) burst number were unaltered by the LFS, paired t-tests p>0.05, n=6 (the average of two sessions per mouse as grey lines). Mean with SEM presented as error bars. (H) Epileptiform bursts, including LL, ML, and HL, were significantly shortened by LFS compared to baseline (Kolmogorov–Smirnov test, p<0.001).

## Discussion

In this study, we compared the antiepileptic effects of optogenetic LFS on the dentate gate at four different sites: two presynaptic and two postsynaptic targets of the MPP-DGC connection inside and outside the sclerotic hippocampus. LFS of MPP fibers in the sclerotic hippocampus effectively reduced the fraction of time spent in seizure state and decreased the number and duration of epileptiform bursts, while LFS of CaMKIIα-positive neurons in MEC had no impact on these measures. Targeting DGCs in the sclerotic septal or non-sclerotic temporal hippocampus with LFS did not reduce the HL burst ratio or number but shortened the epileptiform bursts. In fact, LFS even increased the burst number in the septal DGC stimulation group. Therefore, in the sclerotic hippocampus, the LFS of presynaptic MPP fibers appears to prevent the initiation of focal seizures, whereas the LFS of postsynaptic DGCs shortens the epileptiform activity but promotes the generation of focal seizures.

Our finding that LFS of the MPP-DGC synapse in the sclerotic hippocampus either suppresses or promotes seizures depending on the pre- or postsynaptic stimulation target is in line with the dentate gate hypothesis posing that seizures occur when the gate collapses. Previously, Krook-Magnuson and colleagues (2015) found that on-demand optogenetic inhibition of DGCs in the sclerotic hippocampus terminated ongoing seizures, whereas activating the same cells increased the likelihood of seizure generalization in ihKA mice [38]. Importantly, they stimulated DGCs at 6.7 Hz, and frequencies between 5-10 Hz have been shown to be pro-convulsive in several studies [29,44]. It is significant that hippocampal seizures can be shortened by optogenetic inhibition [38] as well as low-frequency activation as shown in the present study. DGCs across the ipsilateral hippocampus seem to regulate the seizure duration since the LFS of DGCs in both sclerotic and non-sclerotic hippocampus shortened the epileptiform bursts. In contrast, seizure initiation appears to be specific to the sclerotic part of the ipsilateral hippocampus, since only the septal but not temporal DGC stimulation increased the number of focal seizures in ihKA mice.

The lack of seizure suppression by optogenetic LFS of MEC principal neurons was a surprising result, as other studies have shown the opposite in epilepsy models *in vitro* [53,54] and *in vivo* [55]. Namely, 1 Hz optogenetic LFS of CaMKIIα-expressing neurons shortened and reduced the rate of 4-aminopyridine-induced ictal events in brain slices [54] and hindered the acquisition of kindled seizures in mice [55]. Xu and colleagues (2016) suggested that the antiepileptic effect of entorhinal LFS is mediated by GABAergic interneurons in the hippocampus since (1) the entorhinal LFS mostly excited hippocampal interneurons and inhibited hippocampal principal cells, (2) injection of GABAergic receptor antagonists diminished the antiepileptic effects of LFS, and (3) direct activation of hippocampal interneurons mimicked the effects of entorhinal LFS. One possible explanation to why LFS of CaMKIIα-positive MEC neurons did not have an effect on epileptiform activity in our hands is that chronically epileptic ihKA mice have a severe loss of interneurons in the seizure focus with virtually no GABAergic cells left in DG, CA3 and CA1 [33]. In that case, the question remains as to why the stimulation of the same cells at their terminals (i.e. MPP fiber stimulation) was effective. Another explanation would be the spatial limitation of optogenetic stimulation in the entorhinal cortex: the light might have only reached a subpopulation of entorhinal neurons. The CaMKIIα-expressing neurons in MEC are heterogeneous regarding their projection targets (hippocampus, piriform cortex, contralateral entorhinal cortex) and cellular identity (stellate cells and pyramidal neurons) [56]. Thus, we might have targeted an insufficient number of MEC principal neurons projecting to the septal dentate gyrus. We conclude that it is more reliable to activate fibers and nerve terminals at the target than to stimulate the projection neurons at their origin.

As an alternative to MPP fibers, there are other cell populations in the epileptic network that could be targeted with LFS for seizure control. For example, LFS of the anterior nucleus of the thalamus [57], the midline cerebellum [58], the medial septum [59,60], the ventral hippocampal commissure [61,62], and the subiculum [63] have already shown antiepileptic effects in animal models of TLE. In future studies, it would be informative to apply LFS to CA2 neurons in the seizure focus since these cells survive SE well in ihKA mice [37] and MTLE patients [64]. Preliminary results indicate that both chemogenetic and optogenetic inhibition of CA2 pyramidal neurons reduce seizures in chronically epileptic mice [65]. According to a recent study, the contralateral hippocampus could be involved in seizure generation providing another potential target for LFS.

This study has some limitations and raises open questions that remain for future studies. Firstly, it was shown recently that the CaMKIIα-promoter is not only driving ChR2 expression in principal neurons but also in GABAergic interneurons [67], thus we potentially also activated a sub-population of MEC interneurons. Secondly, the use of wire electrodes restricts the interpretation of stimulation responses and their spread. In the future, recordings with silicon probes would aid a better understanding of the presynaptic effects of the LFS and propagation patterns of the stimulus-response in the network. Thirdly, the LFS durations were different in the reanalyzed data with MPP stimulation (one-hour stimulation) and the other LFS targets (three-hour stimulation) as the latter was initially intended for comparison with electric LFS [68]. However, we are confident that the acute seizure-suppressive effects of LFS are not different in one versus three hours. Finally, this study investigated short-term LFS protocols, limiting our interpretation of antiepileptic effects on non-convulsive focal seizures. The effect of LFS on convulsive generalized seizures, which are relatively rare in MTLE, would be a topic of a study with long-term stimulation including both sexes and assessment of cognitive performance. To increase clinical translatability, we would further investigate long-term electric LFS at the sclerotic hippocampus, which has similar seizure-suppressive effects as the optogenetic LFS of MPP [44,68].

In conclusion, presynaptic stimulation of the MPP-DGC synapse within the seizure focus appears critical for the LFS-mediated seizure suppression in the mouse model of MTLE with HS. This effect might be mediated by LFS-induced presynaptic plasticity changes, but the exact mechanism requires further research. Our study provides valuable insights into the potential of LFS as a therapeutic strategy for MTLE and underscores the importance of appropriate target selection for neuromodulation.

## Author contributions

**Piret Kleis**: Conceptualization; Data curation; Formal analysis; Investigation; Project administration; Visualization; Roles/Writing - original draft; **Enya Paschen**: Conceptualization; Data curation; Project administration; Investigation; Methodology; Writing - review & editing; **Ute Häussler**: Methodology; Writing – review & editing; **Carola A. Haas**: Conceptualization; Funding acquisition; Project administration; Resources; Supervision; Writing - review & editing

## Acknowledgments

We thank Jessica Link and Vatsal Jariwala for experimental assistance and data management, Andrea Djie-Maletz for excellent technical assistance and Katharina Heining for her support with the algorithm for epileptiform activity detection and classification.

## Abbreviations

CaMKIIα: Ca^2+^/calmodulin-dependent protein kinase II α

CEMT: Center for Experimental Models and Transgenic Service

cHC: contralateral hippocampus

DBS: deep brain stimulation

DGC: dentate granule cells

HFS: high frequency stimulation

HL: high-load

HS: hippocampal sclerosis

iHC: ipsilateral hippocampus

ihKA: intrahippocampal kainate

LFS: low frequency stimulation

KA: kainate

LFP: local field potential

LL: low-load

ML: medium load

MEC: medial entorhinal cortex

MPP: medial perforant path

MTLE: mesial temporal lobe epilepsy

Rbp4: retinol binding protein 4

RM: repeated measures

SE: status epilepticus

## Appendix

**Table A1.**
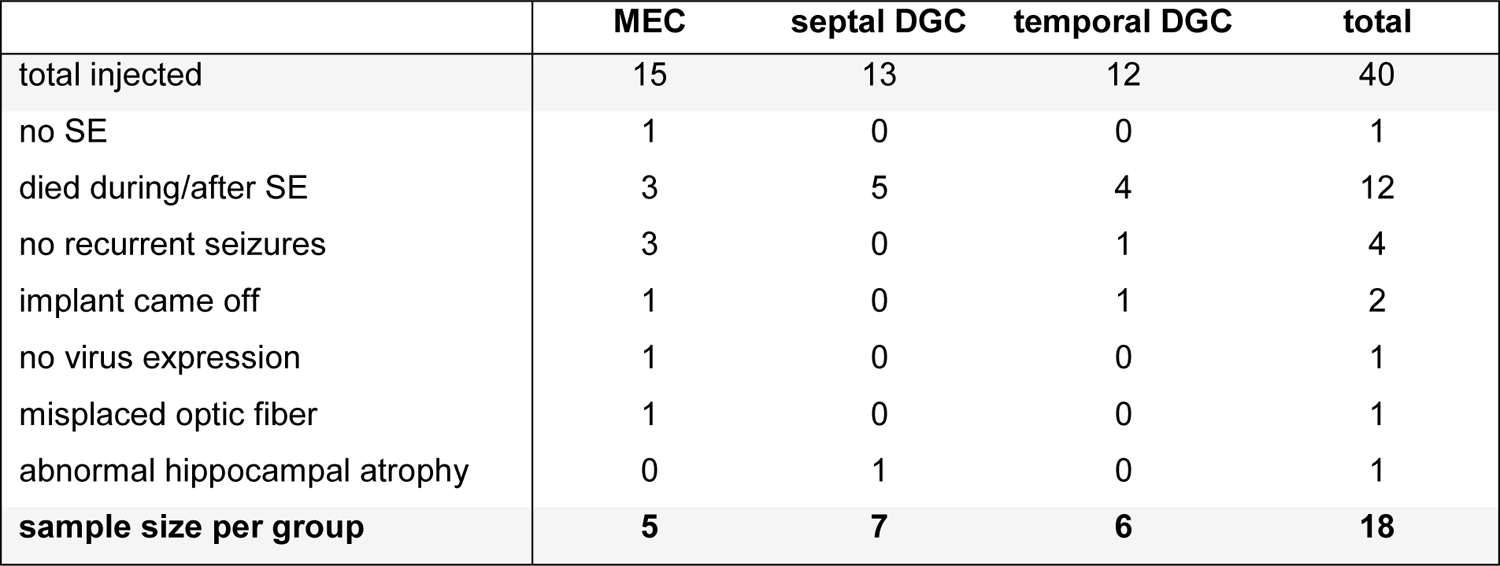
Overview of the animals used for this study and the exclusion criteria.

**Figure A1.**
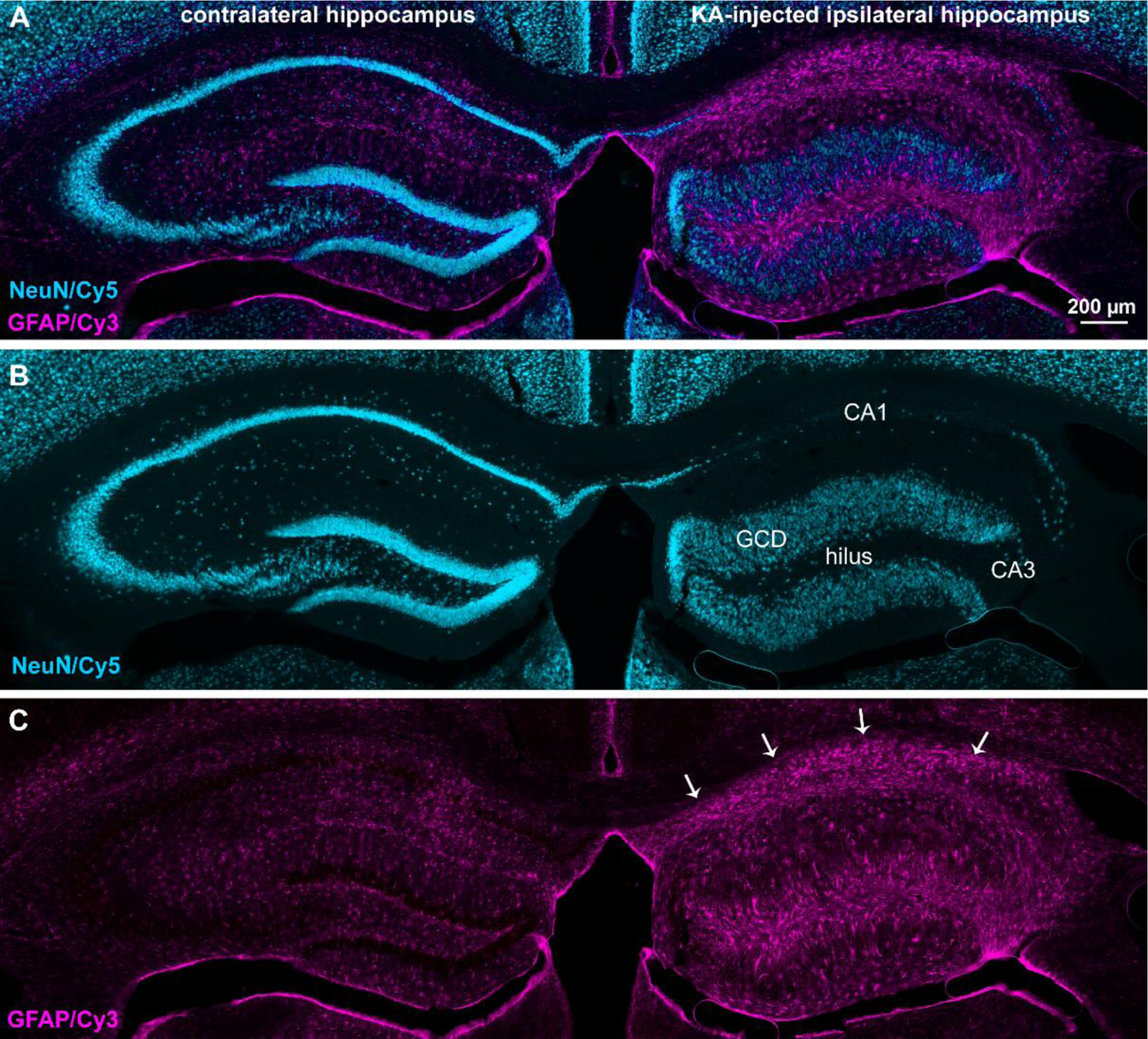
Histological validation of hippocampal sclerosis in the KA-injected hippocampus. (A) A representative image of a coronal section with double-labelling of neurons (NeuN) and astrocytes (GFAP). (B) In the NeuN-labelled section, HS-related changes are clearly visible such as strong neoronal loss in hilus, CA3 and CA1 and granule cell dispersion (GCD). (C) GFAP-labelling demonstrates strong astrogliosis in the KA-injected hippocampus (white arrows).

