## Appendix: Table A1 and Figure A1 for "Low frequency stimulation for seizure suppression: identification of optimal targets in the entorhinal-hippocampal circuit"

### Appendix 1

*Table A1. Overview of the animals used for this study and the exclusion criteria.*

| | MEC | septal DGC | temporal DGC | total |
|---|---|---|---|---|
| total injected | 15 | 13 | 12 | 40 |
| no SE | 1 | 0 | 0 | 1 |
| died during/after SE | 3 | 5 | 4 | 12 |
| no recurrent seizures | 3 | 0 | 1 | 4 |
| implant came off | 1 | 0 | 1 | 2 |
| no virus expression | 1 | 0 | 0 | 1 |
| misplaced optic fiber | 1 | 0 | 0 | 1 |
| abnormal hippocampal atrophy | 0 | 1 | 0 | 1 |
| **sample size per group** | **5** | **7** | **6** | **18** |

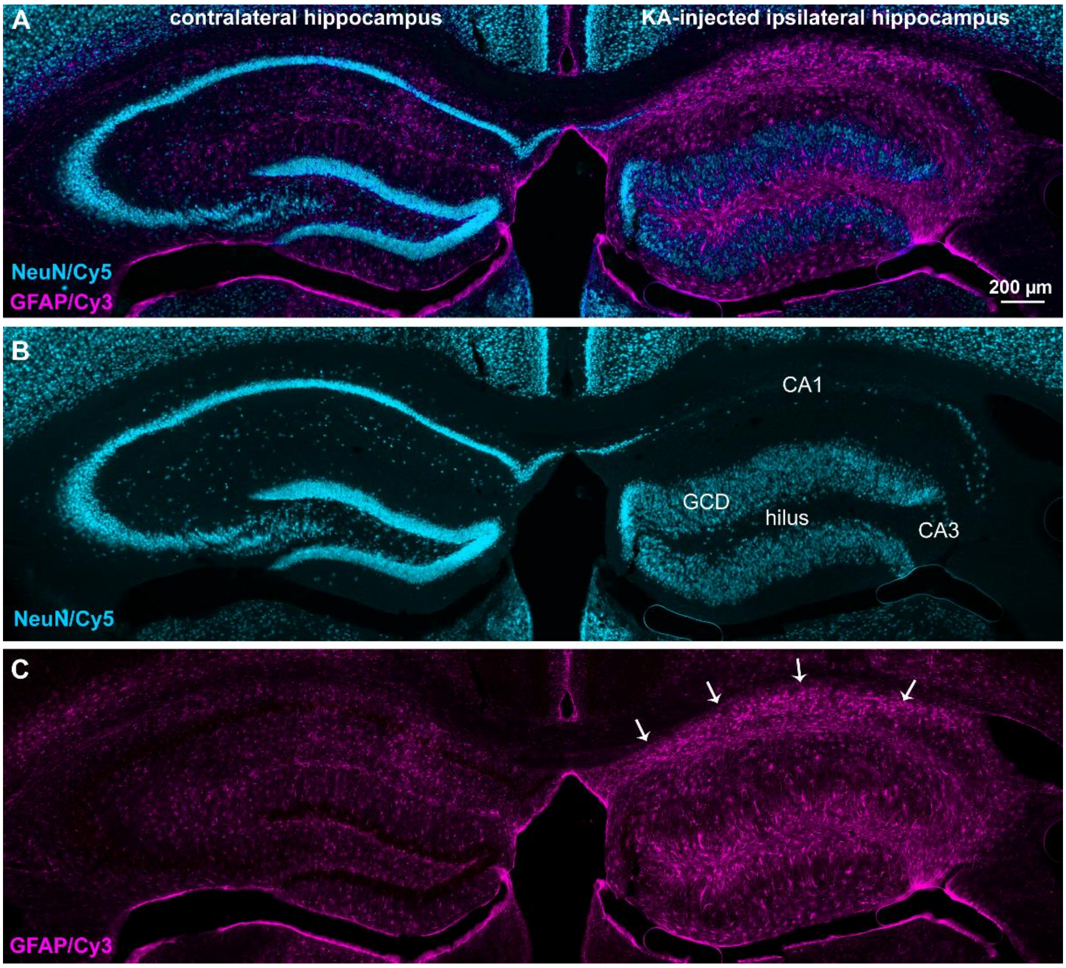


**Figure A1. Histological validation of hippocampal sclerosis in the KA-injected hippocampus.** (A) A representative image of a coronal section with double-labelling of neurons (NeuN) and astrocytes (GFAP). (B) In the NeuN-labelled section, HS-related changes are clearly visible such as strong neoronal loss in hilus, CA3 and CA1 and granule cell dispersion (GCD). (C) GFAP-labelling demonstrates strong astrogliosis in the KA-injected hippocampus (white arrows).
